# Hot and sour: parasite adaptations to honey bee body temperature and pH

**DOI:** 10.1101/2021.07.03.447385

**Authors:** Evan C Palmer-Young, Thomas R Raffel, Jay D Evans

## Abstract

Host temperature and gut chemistry can shape resistance to parasite infection. Heat and acidity can limit trypanosomatid infection in warm-blooded hosts, and could shape infection resistance in insects as well. The colony-level endothermy and acidic guts of social bees provide unique opportunities to study how temperature and acidity shape insect-parasite associations. We compared temperature and pH tolerance between three trypanosomatid parasites from social bees and a related trypanosomatid from poikilothermic mosquitoes, which have alkaline guts.

Relative to the mosquito parasites, all three bee parasites had higher heat tolerance that reflected levels of endothermy in hosts. Heat tolerance of the honey bee parasite *Crithidia mellificae* was exceptional for its genus, implicating honey bee endothermy as a filter of parasite establishment. The lesser heat tolerance of the emerging *Lotmaria passim* suggests possible spillover from a less endothermic host. Whereas both honey bee parasites tolerated the acidic pH’s found in bee intestines, mosquito parasites tolerated the alkaline conditions found in mosquito midguts, suggesting that both gut pH and temperature could structure host-parasite specificity. Elucidating how host temperature and gut pH affect infection—and corresponding parasite adaptations to these factors—could help explain trypanosomatids’ distribution among insects and invasion of mammals.

## INTRODUCTION

Infection by parasites depends on their ability to survive and proliferate under the conditions found in their hosts [1]. Two defining characteristics of this environment are temperature and pH. Host body temperature can profoundly affect host-parasite interactions [2]. In particular, elevated host body temperature due to physiological or behavioral fever limits parasite growth and reduces infection-related morbidity in diverse animals, including insects [3–5]. pH is another driver of microbial establishment [6]. Gut pH contributes to sterilization of food and limits proliferation of opportunistic pathogens [7,8], shaping species-specific resistance to parasites in the insect gut [9]. Knowledge of how host temperature and pH affect host specificity of insect parasites could help to identify host niches and parasite adaptations that affect infection of beneficial insects and potential for insect-vectored, zoonotic spillover to warm-blooded mammals.

The trypanosomatid gut parasites of insects infect a diverse range of hosts—comprising a variety of thermal niches and gut physiologies—with apparently loose host-parasite specificity that remains incompletely understood [10]. The invasion of mammals by a subset of these insect-associated species— the *Leishmania* and *Trypanosoma*—is thought to be limited by mammals’ high body temperatures [11], which can confine infections to (cooler) peripheral body sites even in established mammalian pathogens [12]. In *Leishmania*, where the mammalian stage is intracellular, the low pH of the phagocyte lysosome poses an additional barrier to infection [12]. Nevertheless, putatively monoxenous (i.e., insect-restricted) parasites in the Leishmaniinae sub-family occasionally infect humans [13,14]; such candidate dixenous (i.e., two-host) strains were found in retrospect to be heat-tolerant [13,15]. If temperature and pH limit the establishment of insect trypanosomatids in mammals, these same factors—which vary widely across insect geographic ranges and nutritional niches [16]—could affect the host specificity of parasites among insects as well.

The social honey and bumble bees offer unique opportunities to study parallel adaptations of trypanosomatids to high temperature and low pH in monoxenous trypanosomatids. Whereas most solitary insects have a small body size and limited ability to thermoregulate, social bees inhabit large, thermoregulated colonies with temperatures resembling those of warm-blooded mammals [17,18]. Such high temperatures increase resistance to other pathogens [19,20], and could limit infection by heat-intolerant trypanosomatids as well. Second, bee diets consist of sugar-rich nectar and polysaccharide-rich pollen, which are fermented to organic acids by characteristic gut symbionts that maintain an acidic pH in the honey bee hindgut and rectum [21,22]. This contrasts with the guts of hematophagous Dipteran insects—including mosquitoes—which obtain nitrogen from low-polysaccharide animal blood and have near-neutral to alkaline gut environments [23–25].

To test whether host thermoregulation and diet-associated gut pH can function as filters of trypanosomatid infection in insects, we compared the effects of temperature and pH on growth of phylogenetically related hindgut parasites from honey bees (*Crithidia mellificae* and *Lotmaria passim*), bumble bees (four strains of *Crithidia bombi*, using previously published data [26,27]), and mosquitoes (two strains of *Crithidia fasciculata* [28]). The two major honey bee trypanosomatids—*C. mellificae* [29] and the emerging parasite *Lotmaria passim*, both in the Leishmaniinae [30]—have a global distribution, can reach >90% prevalence in managed colonies, and have been associated with colony collapse on three continents [31–35]. Both species—as well as the bumble parasite *C. bombi* [36]—establish in the hindgut and rectum, the most acidic regions of the intestine [21,37]. Based on the thermal strategies of their host species, we predicted that parasites of highly endothermic honey bees would have greater heat tolerance than parasites from mosquitoes, with intermediate heat tolerance in parasites of bumble bees—which thermoregulate their nests at lower temperatures than do honey bees [38]. We also predicted that parasites of pollen-eating bees would have greater tolerance to acidity than would parasites of blood-consuming mosquitoes, reflecting differences in the diets and gut pH’s of their hosts.

## MATERIALS AND METHODS

### Cell Cultures

*Crithidia mellificae* (ATCC 30254 [29]), *L. passim* (strain BRL [30]) and *C. fasciculata* strains “CFC1” [39] and “Wallace” (ATCC 12857) were obtained from the American Type Culture Collection and collaborators. Honey bee parasites were grown in ‘FPFB’ medium including 10% heat-inactivated fetal bovine serum (pH 5.9-6.0 [40]). Mosquito parasites were grown in brain-heart infusion broth with 20 ug/mL hemin (pH 7.4). All parasites were incubated at 20 °C in vented cell culture flasks and transferred to fresh media every 2 d.

### Temperature experiments

Parasite growth rates were measured by optical density (OD_600_) at temperatures between 20 and 41°C (intervals of 2°C between 23°C and 31°C) on a temperature-controlled microplate reader with 0.1°C resolution (Biotek ‘Synergy’ H1). Cultures were diluted in fresh media to a net OD of 0.040 and aliquoted to 96-well plates containing 120 μL media per well. Measurements were taken every 5 min for 24 h, with 30 s shaking before each read. Each single-temperature block consisted of one 96-well plate with 15 wells (treated as technical replicates) of each of the four parasite strains and 6 cell-free control wells—containing an equal volume of media without parasites—to control for growth-independent changes in OD during incubation. At least two full blocks were conducted at each temperature, to avoid confounding the effects of experimental block and temperature treatment.

### pH experiments

Parasite growth rates were measured between pH 2.1 and 11.3. Aliquots of the base medium for each parasite were first acidified and alkalized to extreme pH levels that inhibited growth in preliminary trials. Treatments were prepared by combining acidified and alkalized media in varying proportions to generate 12 treatments spanning a broad pH range. To initiate the assay, a 12x suspension of cultured cells was added to each treatment for a starting OD of 0.020 in a volume of 120 μL. Each experimental block contained one well per strain plus two cell-free controls of each pH treatment. Growth rates were measured at 29°C for 24 h at 5 min intervals using a microplate reader. Final pH (after addition of fresh media to 1/12 of the final volume) was measured for each treatment using a pH electrode, calibrated immediately prior to measurement. The entire experiment was performed twice, with a slightly narrower pH range in the second block to obtain more complete pH performance curves.

### Comparisons with previous results

To compare thermal performance curves of honey bee parasites and their hosts, we used data for the temperature dependence of force generation during honey bee flight [41] **(Supplementary Fig. 1)**. For comparison to parasites from hosts with intermediate levels of thermoregulation, we used previously published data for thermal performance of four strains of the bumble bee parasite *C. bombi*. For these datasets, growth rates of four strains were measured across temperatures from 17 to 42°C [26], and growth rates of one strain were measured across pH values from 5.0 to 6.2 [27] (**Supplementary Fig. 2**).

### Statistical Analysis

Analyses were conducted using R for Windows v4.0.3 [42]. Models were fit using package “rTPC” [43]. Figures were made with packages “ggplot2” and “cowplot” [44,45]

#### Growth rates

Net OD was calculated by subtracting the average OD from cell-free controls of the corresponding media, treatment, and time point. Growth rates for each well were calculated as the maximum slope of the curve of ln(OD) vs. time, obtained by fitting a rolling linear regression to each 4 h window of the growth curve [46]. The first 2 h of each run were excluded to allow OD readings to stabilize. We used only slopes with r^2^ values of >0.95 and >0.90 for the temperature and pH experiments, respectively, and assigned a growth rate of zero to samples where the average slope of the growth curve was negative. For temperature experiments, we used the median growth rate among the 15 replicates within each block, to avoid pseudoreplication within each implementation of the temperature treatment [47].

#### Temperature models

We modeled the temperature dependence of growth for each trypanosomatid strain using a Sharpe-Schoolfield equation modified for high temperatures [46,48,49].

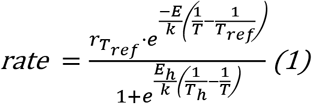

In Equation (1), *rate* refers to the maximum specific growth rate (in h^-1^); *r_Tref_* is the growth rate (in h^-1^) at an arbitrary calibration temperature *T_ref_* (fixed at 20°C); *E* is the activation energy (in eV), which primarily affects the upward slope of the thermal performance curve (i.e., sensitivity of growth to temperature) at suboptimal temperatures; *k* is Boltmann’s constant (8.62·10^-5^ eV·K^-1^); *E_h_* is the deactivation energy (in eV), which determines how rapidly the thermal performance curve decreases at temperatures above *T_pk_; T_h_* is the high temperature (in K) at which growth rate is reduced by 50% (relative to the value predicted by the Arrhenius equation—which assumes a monotonic, temperature-dependent increase) [49]; and *T* is the experimental incubation temperature (in K).

#### pH models

To describe the effects of pH on growth rates, we used a biphasic logistic model that describes sigmoidal decreases in growth rate at low and high pH.

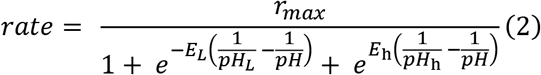

In Equation (2), *r_max_* is the specific growth rate at the optimum pH; *E_L_* and *E_h_* correspond to the rates of deactivation at low and high pH, respectively; and *pH_L_* and *pH_h_* represent the pH values at which growth rate is reduced by 50% relative to *r_max_*. Due to absence of high-pH measurements for *C. bombi*, models for this species were fit using a standard (monophasic) logistic regression, which omitted the second term of the denominator in Equation (2).

Models were optimized using nonlinear least squares, implemented with R packages rTPC and nls.mulstart [43]. Confidence intervals on parameter values and predicted growth rates were obtained by bootstrap resampling of the residuals (10,000 model iterations) [50]. We also used the bootstrap model predictions to estimate the following traits: temperatures of peak growth rate (*T_pk_*) and 50% inhibition relative to the peak value (*IT_50_*); pH of peak growth (*pH_pk_*); and pH niche breadth (i.e., the number of pH units between *pH_L_* and *pH_h_*). The 0.025 and 0.975 quantiles for parameter estimates, predicted growth rates at each temperature, and traits derived from bootstrap predictions were used to define 95% confidence intervals. Strains were considered significantly different from each other when their 95% confidence intervals did not overlap.

## RESULTS

### Temperature experiments

Thermal performance curves **(Fig. 1)** and model parameters **(Fig. 2)** showed higher heat tolerance in the two honey bee parasites than in the mosquito parasites. *Crithidia mellificae* (peak (*T_pk_*): 35.42°C, 50% inhibition (*IT_50_*): 38.7°C) grew well throughout the temperature range found in honey bee hives during brood-rearing (33.8-37°C [18]) and exhibited the peak growth temperature closest to that of *A. mellifera* (38.4°C [41]; **Supplementary Fig. 1**).The heat tolerance of *L. passim* (*T_pk_*: 33.47°C, *IT_50_*: 36.97°C) was approximately 2°C less than that of *C. mellificae*, with predicted growth rates reduced by >50% at the upper end of the thermal range found in colonies **(Fig. 1)**. Thermal performance curves and parameter estimates were similar for the two strains of *C. fasciculata*, where temperatures of peak growth (strain CFC1: 30.92°C, strain Wallace: 31.58°C) and 50% inhibition (CFC1: 35.27°C, Wallace: 35.48°C) were approximately 2°C lower than for *L. passim* and 4°C lower than for *C. mellificae* **(Fig. 2)**. Nevertheless, both strains had peak growth temperatures (*T_pk_*) that exceeded the mean *T_pk_* for a variety of traits in diverse mosquito species (28.4°C [51], **Fig. 2**).

**Figure 1.**
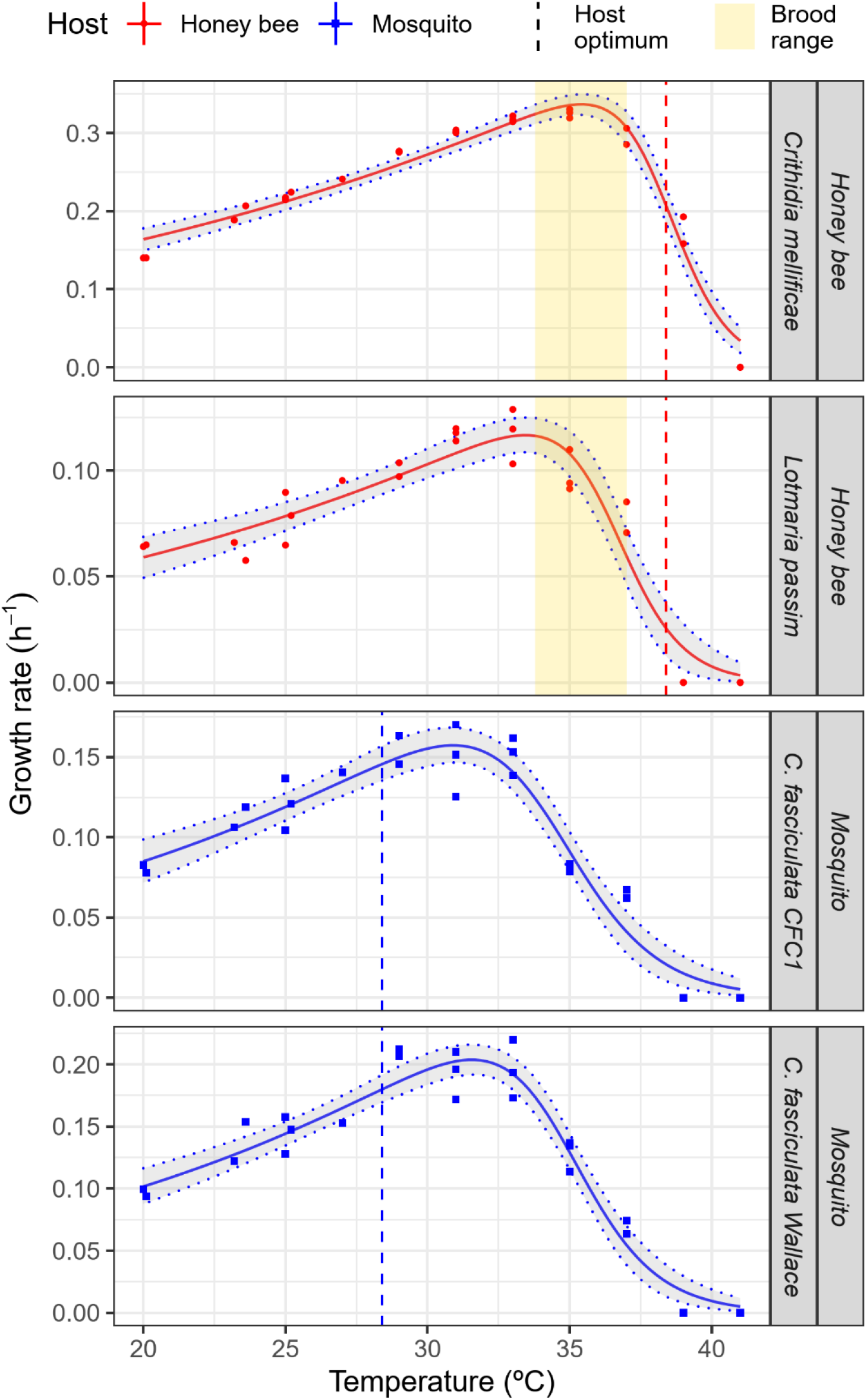
Thermal performance curves for trypanosomatid parasites from honey bees (*Crithidia mellificae, Lotmaria passim*) and mosquitoes (*Crithidia fasciculata*). The right-most facet label indicates the strain’s host of origin. Each point represents the median specific growth rate (h^-1^) from one 15-replicate experiment, with color and shape corresponding to the parasite’s host. Lines and shaded bands show predictions and 95% bootstrap confidence intervals from Sharpe-Schoolfield models [43,49]. Vertical lines show optimum temperatures for honey bees (estimated from force production during flight [41]) and mosquitoes (mean of 88 traits [51]). Vertical band (in yellow) shows temperature range for honey bee brood incubation [18]. See Supplementary Figure 1 for full thermal performance curve of honey bee force production.

**Figure 2.**
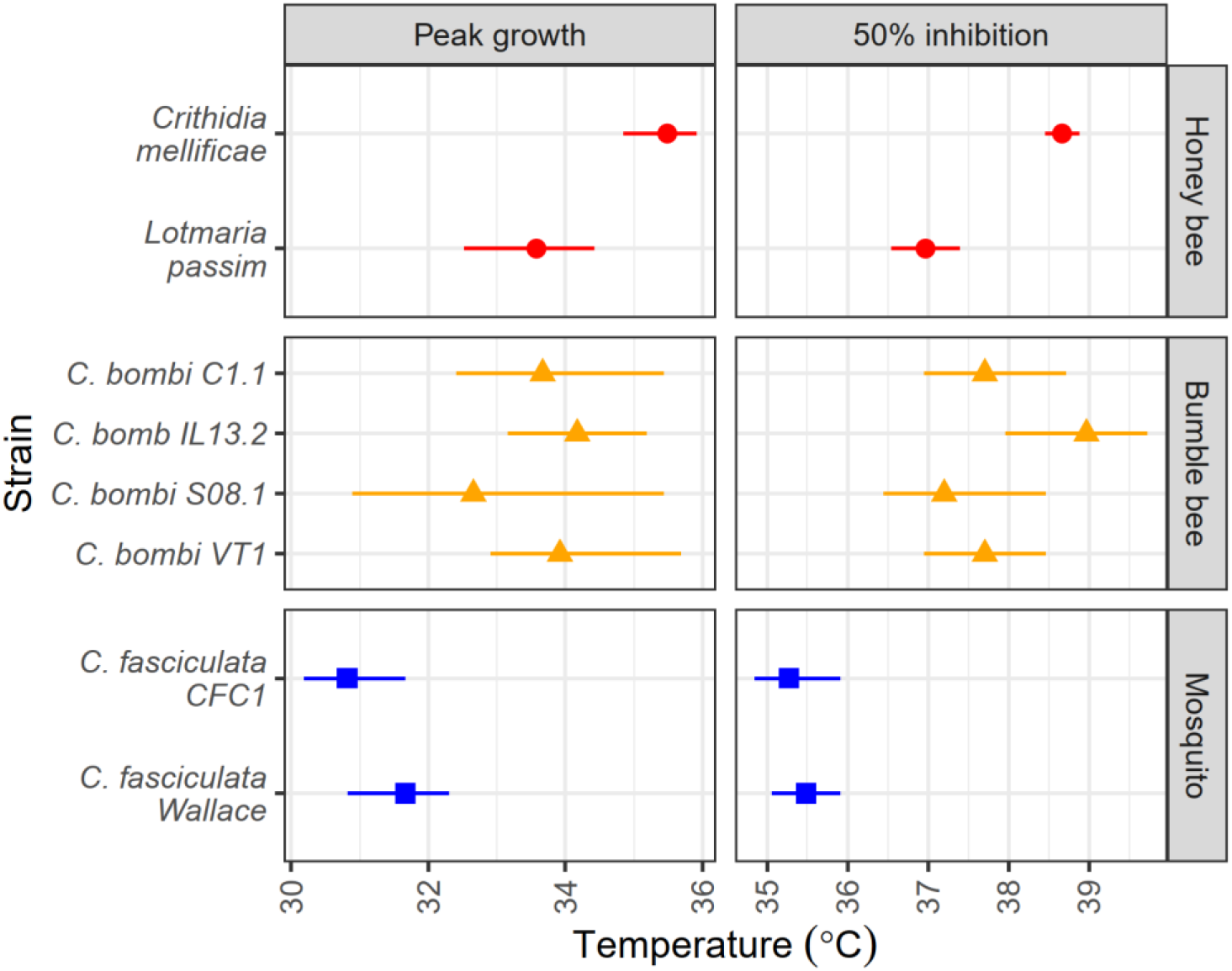
**Temperatures of peak growth and 50% inhibition of growth rate** for parasites of honey bees (*Crithidia mellificae, Lotmaria passim*), bumble bees (*C. bombi*, tested in [26]), and mosquitoes (*C. fasciculata)*. Points and error bars show estimates and 95% bootstrap confidence intervals for predictions from Sharpe-Schoolfield models. See Supplementary Figure 2 for full thermal performance curves for *C. bombi*. Estimates for additional model parameters are shown in Supplementary Figure 3.

Thermal performance curves of *C. bombi* from bumble bees (*T_pk_*: 33.67°C; *IT_50_*: 37.90°C, **Fig. 2, Supplementary Fig. 2)** most resembled that of *L. passim*. Although the coarser 5°C temperature interval for the published *C. bombi* data resulted in higher uncertainty, all four strains of this species appeared to have at least 2°C higher inhibitory temperatures (*IT_50_*) than did *C. fasciculata* **(Fig. 2)**. Activation energies (*E*) ranged from 0.39 eV (*C. mellificae*) to 0.52 eV (*C. fasciculata* strain Wallace), well within the range observed across a diversity of physiological and ecological rates (median 0.55 eV [52], **Supplementary Fig. 3**). High-temperature deactivation energies (*E_h_*) ranged from 5.18 eV (*C. fasciculata* Wallace) to 8.29 eV (*C. mellificae*), consistent with the steep decline at high temperatures that is typical of thermal performance curves [52] **(Supplementary Fig. 3)**.

### pH experiments

We observed the greatest tolerance to acidity in the two parasites of honey bees, each of which grew at nearly two units’ lower pH than either *C. fasciculata* or the previously tested *C. bombi*. Both maintained strong growth at the pH of the honey bee rectum (pH 5.2 [21] **(Fig. 3)**. *Crithidia mellificae* had the broadest pH niche, with the greatest tolerance of both acidity (50% low-pH inhibition (*pH_L_*): 3.07, 95% CI: 2.97-3.25) and alkalinity (50% high-pH inhibition (*pH_h_*): 9.93, CI: 9.55-10.21, **Fig. 4**). *Lotmaria passim* was nearly as tolerant of acidity as was *C. mellificae* (*pH_L_*: 3.44, CI: 3.35-3.53) but grew weakly above pH 7 (*pH_h_*: 7.33, CI: 7.24-7.43), with peak growth pH (5.57, CI: 5.20-5.76) closely matched to that of the host rectum **(Fig.’s 3-4)**.

**Figure 3.**
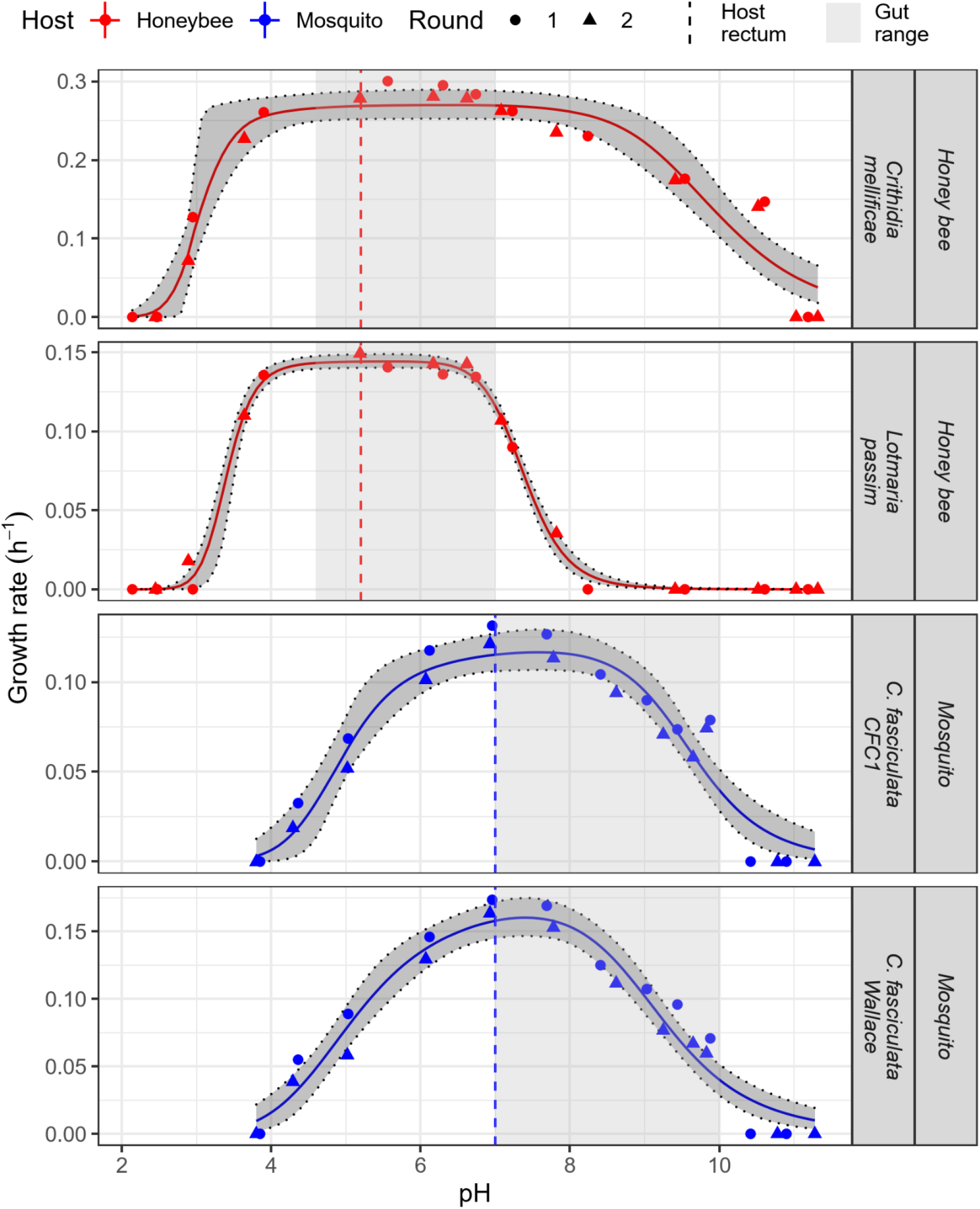
**Effects of pH on growth** of trypanosomatid parasites from honey bees (*Crithidia mellificae, Lotmaria passim*) and mosquitoes (*Crithidia fasciculata*). The right-most facet label indicates the strain’s host of origin. Each point represents the specific growth rate (h^-1^)) from one sample. The experiment was conducted over two experimental blocks (Round 1: circles; Round 2: triangles). Lines and shaded bands show predictions and 95% bootstrap confidence intervals from biphasic logistic models. Vertical lines and shaded regions show pH of the rectum (primary site of parasite infection) and range of the gut overall, as measured previously in honey bees [21,37] and *Culex* mosquitoes [25].

**Figure 4.**
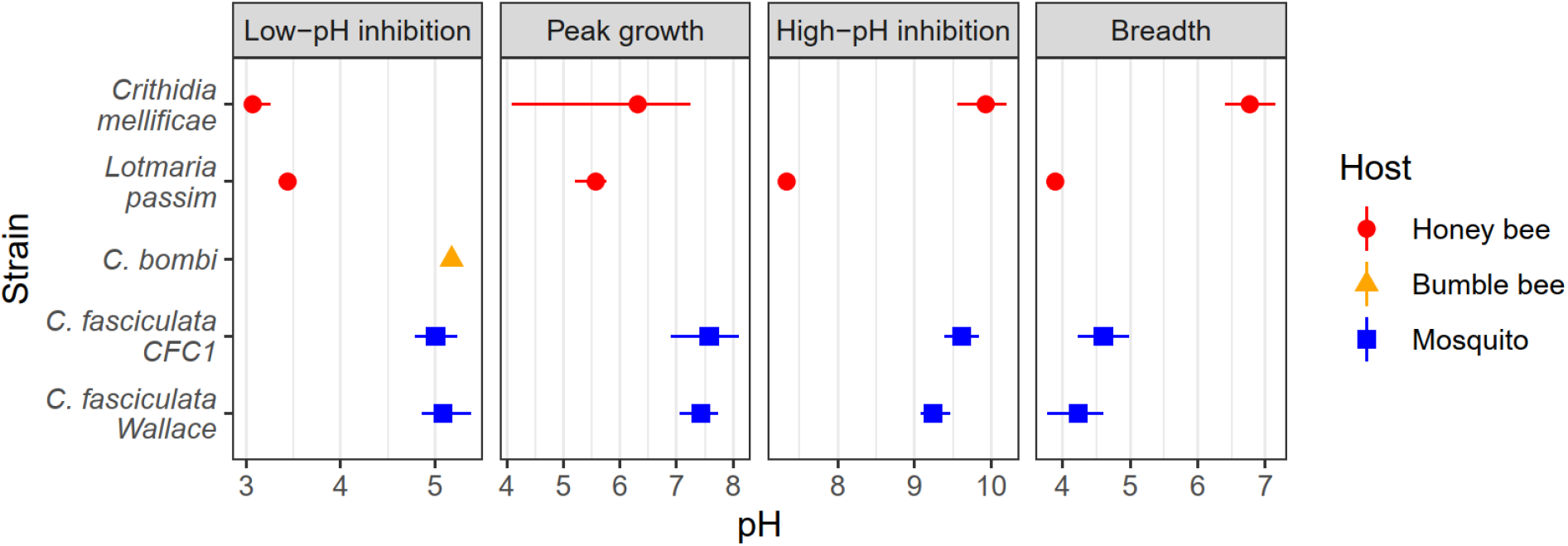
Estimates for pH of peak growth, 50% inhibition of growth rate due to low and high pH, and pH niche breadth. (i.e., difference between estimates of 50% inhibition due to low and high pH) for parasites of honey bees (*Crithidia mellificae, Lotmaria passim*), bumble bees (*C. bombi* strain IL13.2, tested in [27]), and mosquitoes (*C. fasciculata*). Points and error bars show estimates and 95% bootstrap confidence intervals for predictions from biphasic logistic models. Colors and shapes correspond to host of origin. See Supplementary Figure 4 for full model predictions for *C. bombi*. Estimates for additional model parameters are shown in Supplementary Figure 5.

In contrast, both strains of *C. fasciculata* grew fastest at neutral to weakly basic pH (*pH_pk_* for CFC1: estimate 7.58, CI: 6.90-8.10; Wallace: estimate 7.42, CI: 7.05-7.73, **Fig.’s 3-4**). Although tolerance of acidity was not as great as in the honey bee parasites (*pH_L_* for CFC1: 5.01, CI: 4.71-5.24; Wallace: 5.08, CI: 4.86-5.39), the two strains were tolerant of alkaline conditions (*pH_h_* for CFC1: 9.62, CI: 9.39-9.84; Wallace: 9.24, CI: 9.01-9.47) that approached those found in the midgut of their host *Culex pipiens* [25] **(Fig.’s 3-4)**. Acidity tolerance in *C. bombi* (*pH_L_* 5.18, CI: 5.17-5.19) was indistinguishable from that of *C. fasciculata* (**Fig. 4**; see **Supplementary Fig. 4** for full *C. bombi* curves). *Crithidia bombi* was also notable for its steep decline in growth rate between pH 6 and pH 5 [27], which was reflected in an estimate for deactivation energy (parameter *E_l_*) more than 6-fold higher than that of the strains tested here **(Supplementary Fig. 5)**.

## DISCUSSION

Our results indicate the importance of colony-scale endothermy in social bees as a filter for gut parasites. All the parasites from endothermic social bees showed greater heat tolerance than did parasites from mosquitoes. This was particularly notable for *C. mellificae*, which exhibited superior heat tolerance to all previously studied, poikilothermic tropical insect-associated trypanosomatids noted for their heat tolerance. For example, growth of *Crithida luciliae thermophila* (since renamed *C. thermophila* [53]), *Crithidia hutneri* [54], and *Leptomonas pessoai* (renamed *Herpetomonas samuelpessoai* [55,56]) all grew faster at 28°C than at 37°C. Growth of *Leptomonas seymouri*—which occasionally infects humans [14]—was likewise poor at 37°C [57]. In contrast, growth of our *C. mellificae* was approximately 30% faster at 37°C than at 28°C. Such heat tolerance was suggested by Cosgrove and Mcghee [58], whose review stated that an unnamed trypanosomatid from *Vespula squamosa* (presumably ATCC strain 30862 of *C. mellificae*) grew in avian embryos at 37°C with no prior acclimation. However, the relevant reference [59] did not mention *C. mellificae*. Of note, the species that was shown to maintain strong growth in embryos at 37° was *Crithidia acanthocephali* [60]. Although originally isolated from a Hemipteran [60], sequences matching this species were recently amplified from honey bees in Spain [61]; the parasite’s heat tolerance could facilitate its survival in bees.

The warm-blooded mammal-like temperatures of a breeding honey bee colony [18] likely preclude infection by trypanosomatids with low heat tolerance, and could exert positive selection for heat tolerance within parasite lineages. For parasites that do establish in colonies, our results suggest that high colony temperatures might reduce infection intensities. Even growth of the most heat-tolerant parasite (*C. mellificae*) peaked at a lower temperature than did flight performance of honey bee hosts (38.4°C, **Fig. 1**). Peak performance temperatures of flight muscle [62] and respiration [63] in bumble bees are also high (>40°C). This suggests that increases in temperature could favor increases in host metabolic performance—perhaps including immune function—while inhibiting parasite growth. Honey and bumble bee gut symbionts—which enhance resistance to *C. bombi* [64]—are likewise heat-tolerant. Honey bee symbionts have standard culturing temperatures of 35-37°C [65], can grow at temperatures up to 44°C [66], and tolerate hour-long heat shock at 52°C [66]. A *Lactobacillus* species from bumble bees was similarly thermophilic, with a peak growth temperature of approximately 40°C [26]. High temperatures could therefore enhance the antiparasitic activities of these symbionts as well as performance of the bee immune system [27], harnessing the bees’ socially enabled thermoregulation and core gut microbiota for defense against infection.

Our results suggest that maintenance of high, ‘social fever’-like colony temperatures would be particularly effective against the relatively heat-susceptible *L. passim* and *C. bombi*. Growth rates of *L. passim* dropped by approximately 50% over the 3.2°C range found in brood-rearing honey bee colonies **(Fig. 1)**. Similarly, infection of *C. bombi* was 81% lower at 37°C than at 21° [67]. Inoculations of honey bees with *C. mellificae* were likewise less successful at 35°C than at 29°C (albeit in separate experiments [29]). Our results also suggest that bees may become increasingly susceptible to infection as they transition from activities at the well-heated colony core to the cooler and more variable periphery, or to foraging outside (at age 10-25 d [68]). Observations of experimentally infected, colony-reared bees— which showed a 10-fold increase in parasite mRNA between ages 7 and 27 d [69]—are consistent with these predictions. However, similar age-related infection dynamics were observed in caged bees at constant temperatures [69], suggesting that other age-related factors could also contribute to this pattern.

Honey bee trypanosomatid infection intensities are inversely related to temperature in field colonies as well [70]. In managed US colonies, *L. passim* infection intensity (originally described as *C. mellificae* [30,71]) peaked in mid-winter, when colony core temperatures average 14°C lower than in summer [18]. Such temperature-dependent infection dynamics could explain the associations between trypanosomatid infection and overwinter colony collapse [32]. Seasonal susceptibility of colonies to infection could be exacerbated by landscape, chemical, and nutritional factors that impair thermoregulation [72,73]. For example, colonies from agricultural areas had average winter temperatures 8°C lower than did colonies from grasslands [74], highlighting how land use changes could affect temperature-mediated resistance to an emerging infectious disease.

*Lotmaria passim’s* low heat tolerance relative to *C. mellificae*, susceptibility to the high temperatures found in honey bee colonies, and apparently recent global emergence in *A. mellifera* [30] all invite speculation of a recent host shift from a less endothermic bee species. The Asian honey bees *Apis cerana* [75] and *A. dorsata* [76] have ~2°C lower brood temperature optima relative to *A. mellifera* [18]—matching the ~2°C difference in optimal and inhibitory temperatures between *C. mellificae* and *L. passim*. *Apis cerana* harbored an *L. passim* haplotype basal to the strains found on other continents [77], providing circumstantial phylogenetic evidence for an Asian parasite origin. Such a host shift could parallel the worldwide dispersal of the now ubiquitous microsporidian *Nosema ceranae* from *A. cerana* [78].

Our findings of acid tolerance in parasites of honey bees and alkaline tolerance in parasites of mosquitoes suggest that gut pH—itself a reflection of diet, digestive physiology, and microbiota—is also an important driver of host specificity in trypanosomatid parasites of insects. The tolerance of acidic conditions shown by honey bee parasites—and the low optimum pH of the emerging parasite *L. passim*—reflect the typically acidic pH found in the honey bee rectum where these parasites establish [21,29,30]. This tolerance of acidity was noted by Langridge and McGhee in their isolations of *C. mellificae* [29]. The honey bee’s low gut pH results from fermentation of pollen polysaccharides by the characteristic bee gut microbiota [21,22]. In humans, acidic intestinal and fecal pH’s likewise reflect the intake and subsequent fermentation of dietary polysaccharides [79], with consequences for microbiome composition and growth of opportunistic pathogens [7,80]. The pH of the bee rectum—which at pH 5.2 is over a full pH unit more acidic than the already pathogen-inhibiting feces of humans consuming fiber-rich vegan diets (pH 6.3 [80])—may likewise provide protection against opportunistic invaders including non-specialist trypanosomatids.

Although standard trypanosomatid culture media is neutral to weakly basic (e.g., brain heart infusion broth, pH 7.4), enhancement of growth under acidic conditions has been reported before. For example, growth of *H. samuelpessoai* occurred between pH 4 and pH 9 [56]. In addition to *C. mellificae*—described as ‘acidophilic’, with optimum growth at pH 5 [29]—McGhee described enhanced growth under acidic conditions (pH 5 vs. pH 8) in three additional trypanosomatids and found growth exclusively at low pH in two others [81]. All these acidophilic species were isolated from hemipteran hosts; two were from the giant milkweed bug *Oncopeltus fasciatus*, whose gut pH (4.6-5.4 [82]) resembles that of honey bees— suggesting potential for bee-hemipteran parasite exchange.

In contrast—and concordant with our results—the parasite species that thrived under basic conditions (including *C. fasciculata*) were from Dipterans [81], where gut pH is typically extremely alkaline. For example, the original host of our *C. fasciculata* (*Culex pipiens*) has a midgut pH greater than 10 in larvae [25]—yet this life stage can still be infected by *C. fasciculata* [28]. Similarly high pH’s occur in the larval guts of other Diptera (e.g., midgut pH of 11 in Bibionid larvae [24]. In mosquito adults, the midgut is near pH 6 in sugar-fed adults [83], but is alkalized to pH 8.5-9.5 following ingestion of blood [23]. Adaptations to these conditions are reflected in our results, with both *C. fasciculata* strains growing fastest near neutral pH (6-8) and remaining viable up to pH 10 **(Fig. 3)**, consistent with previous characterizations [84]. Intriguingly, the difference in pH optima between the honey bee parasite *L. passim* and the mosquito parasite *C. fasciculata* matched almost exactly the differences between the optima for the mammalian tissue (amastigote, pH 5.5) and insect (promastigote, pH 7-7.5) stages of *Leishmania* [12]. This raises the question of whether differences in pH tolerance among species of monoxenous taxa and between life stages of dixenous taxa can be explained by similar mechanisms, and whether tolerance of acidity is correlated with tolerance of high temperature (as in *Leishmania* [12]).

Contrary to predictions, the bumble bee parasite *C. bombi* did not exhibit the high tolerance of acidity found in the honey bee parasites. The single report of bumble bee gut hindgut pH that we could locate (pH 6.25 from *Bombus fervidus* [16]) is substantially higher than the pH <5.2 measured in honey bees [21,37], but a close match to the pH 6.0-6.2 that yields optimal growth of *C. bombi* (**Supplementary Fig. 4**, [27]). Although honey and bumble bees have similar pollen- and nectar-based diets and gut microbial communities [85]—which might be expected to result in similar gut pH—they exhibit marked differences in physiology and behavior. Bumble bees have a more rapid intestinal transit time than do honey bees [86], leaving less time for acid-generating fermentation. In contrast, honey bees not only have slower baseline transit times, but also fastidiously refrain from defecation in the colony—a behavior not exhibited by bumble bees [87]. As honey bees spend the first 10-25 d in the colony before they forage outdoors [68], the pollen-rich rectal contents have considerable time to acidify. During the winter, honey bees commonly retain rectal contents for several months while confined in the colony [88]. Meanwhile, they continue to ingest pollen, with their distended guts exhibiting increases in populations of fermentative hindgut bacteria [89]. We hypothesize that these behaviors result in lower gut pH—and greater selection on parasites for tolerance of acidity—in honey bees than in bumble bees.

The same heat tolerance that allows insect trypanosomatids to infect endothermic bees could also pre-adapt parasites for infection of warm-blooded mammals. Several supposedly monoxenous species have been found in humans—often together with the expected *Leishmania* [13,14,58]—and proven infectious in the glands of opossums and the skin and organs of mice [13,90], demonstrating the ability to proliferate at 37°C. Intriguingly, trypanosomatids with 100% GAPDH sequence identity to *C. mellificae* were recently isolated from the blood of numerous wild mammals in Brazil [91,92]. The viability of these parasites at 37°C [92]-consistent with our findings—would permit survival in the mammalian bloodstream, perhaps additionally aided by parasite acclimation to high temperatures in honey bee colonies. Given that *L. seymouri*—one of the closest known relatives of *C. mellificae* [30]— occasionally infects humans [14] despite minimal growth at 37°C [57], corresponding infection of mammals by *C. mellificae* seems plausible. Although pathways of transmission remain unclear, we have shown that *C. mellificae* from honey bees can proliferate in bees of other families—including halictids, which are attracted to mammalian perspiration [93]. The impressive range of pH tolerance shown here could also support its survival in other, possibly hematophagous hosts with diverse gut physiologies.

## CONCLUSIONS

Our interspecific comparisons—including the first tests of temperature and pH tolerance in the emerging parasite *L. passim*—implicate colony-level endothermy and diet- and microbiome-related changes in gut acidity as drivers of host specificity in insect trypanosomatids. Our results also provide a mechanistic explanation for the relative resistance of honey bees to trypanosomatids from other insects [94] and the recent findings of *C. mellificae*—a presumed monoxenous parasite—in a variety of warm-blooded mammals [91,92]. Escape from parasites could be one factor that favors the evolution of energetically costly social endothermy and maintenance of gut symbiont communities in insects, providing infection-related benefits that parallel those found in homeothermic vertebrates while exerting parallel selective pressures on parasites.

## ACKNOWLEDGMENTS

We thank the ATCC, Michael and Megan Povelones, Stephen Beverley, Ryan Schwarz, and Ben Sadd for parasite strains and culturing advice; Daniel Padfield for R scripts; and anonymous reviewers for their service in improving the manuscript.

## FUNDING

This project was funded by the USDA Agricultural Research Service; a USDA-NIFA Pollinator Health grant to JDE; a North American Pollinator Protection Campaign Honey Bee Health Improvement Project Grant and an Eva Crane Trust Grant to ECPY and JDE; and an NSF-CAREER grant (IOS 1651888) to TRR. Funders had no role in study design, data collection and interpretation, or publication.

## CONFLICTS OF INTEREST

The authors declare that they have no conflicts of interest.

## DATA AVAILABILITY

All data are supplied in the Supplementary Information, Data S1.

## AUTHORS’ CONTRIBUTIONS

ECPY conceived the study. ECPY and TRR designed experiments. ECPY conducted experiments, analyzed data, and drafted the manuscript with guidance from JDE and JDE. All authors revised the manuscript and gave approval for publication.

## MEDIA PROMOTION

High body temperature and acidic gut pH are two factors that inhibit parasitic infection. The high colony temperatures and acidic guts of social bees relative to other insects provide unique opportunities to test how temperature and acidity shape insect-parasite associations and potential for spillover into warm-blooded mammals. We show that parasites of honey bees have greater tolerance of heat and acidity than do related parasites of mosquitoes, which lack both temperature regulation and gut acidity. This suggests that honey bees’ colony-enabled temperature regulation and gut chemistry provide resistance to non-specialist parasites, favoring the same parasite traits needed for mammalian infection.

## Notes

### Competing Interest Statement

The authors have declared no competing interest.

