## Supplementary information for "Hot and sour: parasite adaptations to honey bee body temperature and pH"

Running title: Thermal and pH niches of bee and mosquito parasites

### SUPPLEMENTARY FIGURES

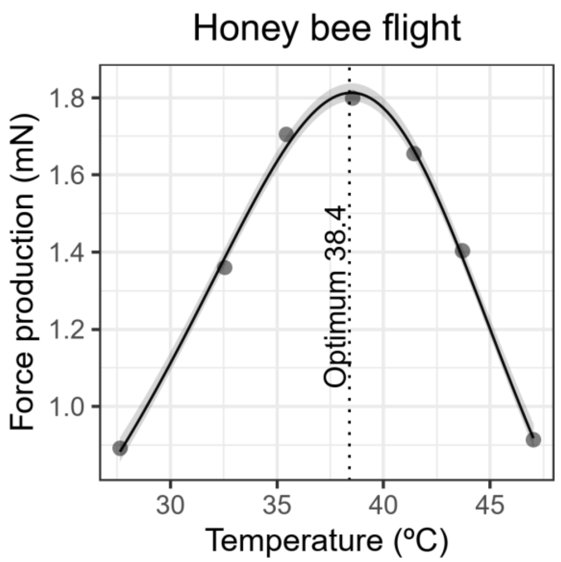

**Supplementary Figure 1. Thermal performance curve for force production by honey bees during tethered flight.** Points show raw data extracted from Harrison and Fewell 2002 [1]. Line and shaded band show predictions and 95% bootstrap confidence intervals from Sharpe-Schoolfield model [2,3].

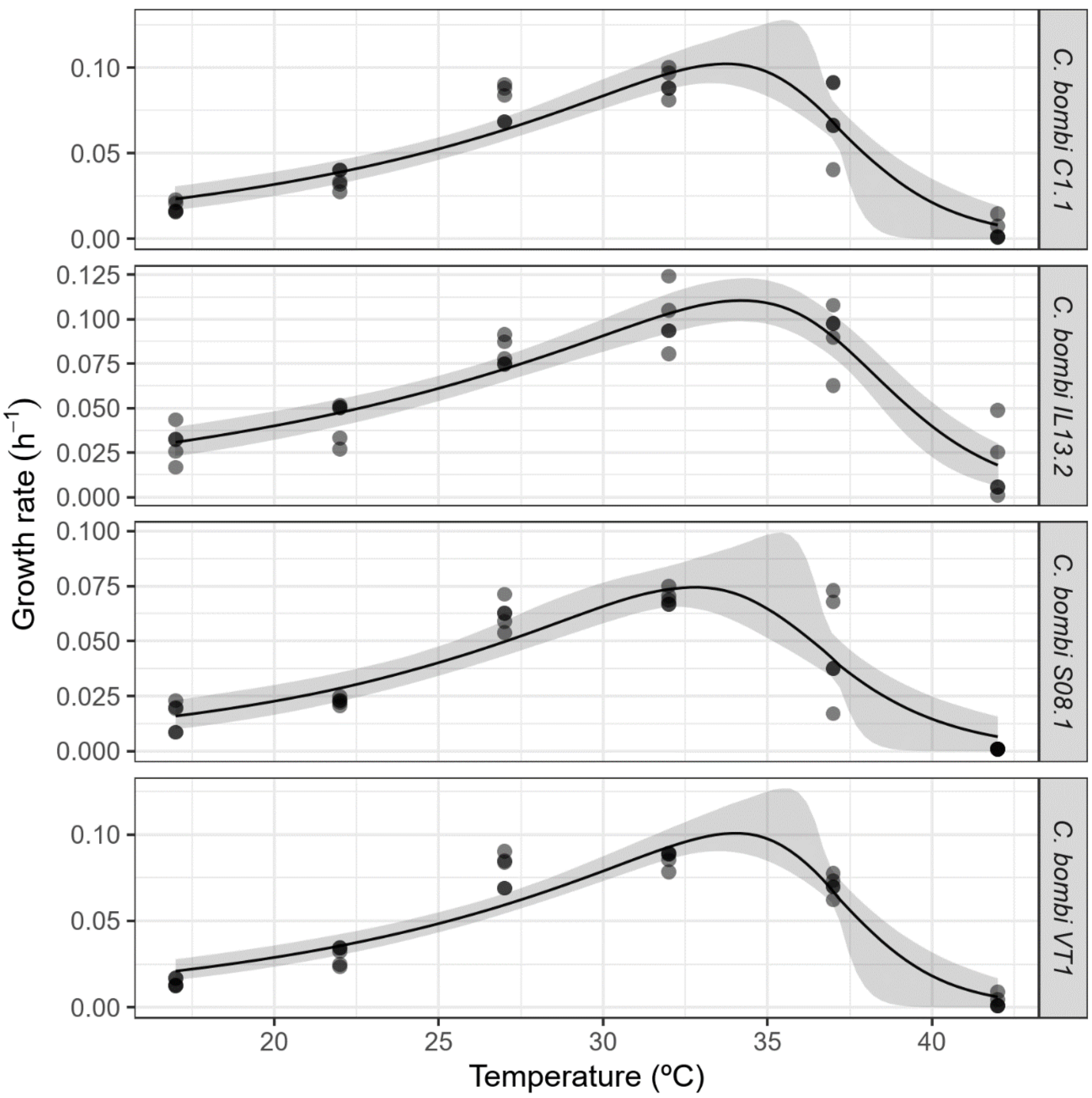

**Supplementary Figure 2. Thermal performance curves for *Crithidia bombi* strains.** Points show raw data. Lines and shaded bands show predictions and 95% bootstrap confidence intervals from Sharpe-Schoolfield model [2,3]. The right-most facet label indicates the strain's host of origin. Data for *C. bombi* are taken from Palmer-Young, Raffel, and McFrederick [4].

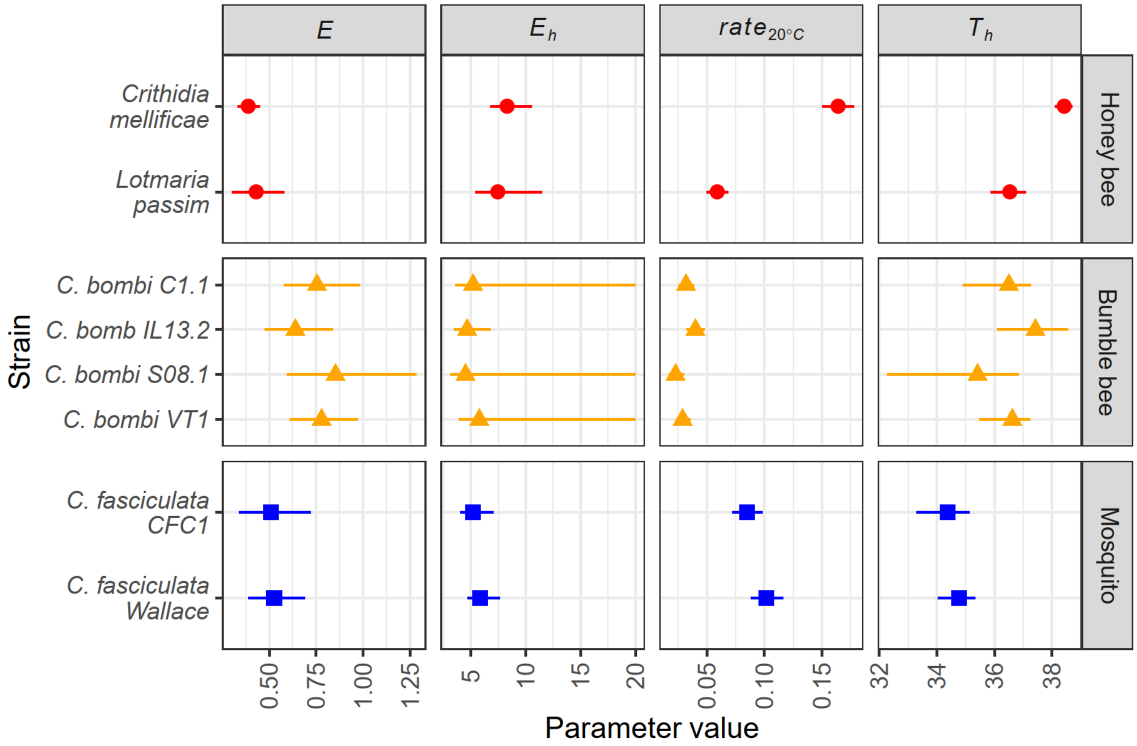

**Supplementary Figure 3. Parameter estimates for models of temperature-dependent growth** by parasites of honey bees (*Crithidia mellificae, Lotmaria passim*), bumble bees (*C. bombi,* tested in [4]), and mosquitoes (*C. fasciculata*). Facets correspond to model parameters (*E:* activation energy (in eV); $E_h$: high-temperature deactivation energy (in eV); $rate_{20°C}$: predicted growth rate at 20 °C (in $h^{-1}$); $T_h$: temperature at which rate is reduced by 50% due to high-temperature deactivation (calculated in K, shown in °C)). Points and error bars show estimates and 95% bootstrap confidence intervals for predictions from Sharpe-Schoolfield models. Colors and shapes correspond to host of origin. See Supplementary Figure 2 for full model predictions for *C. bombi.*

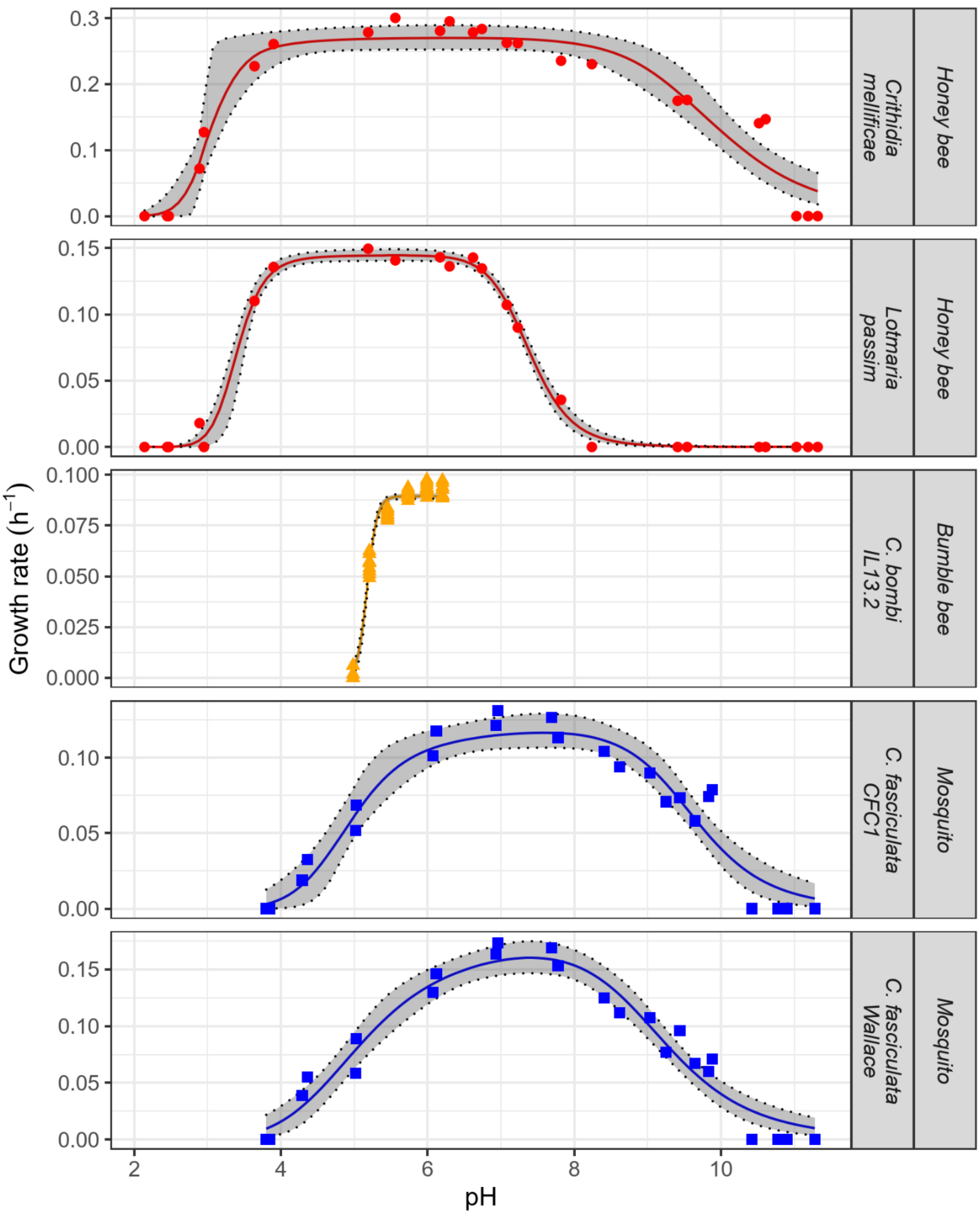

**Supplementary Figure 4. Effects of pH on growth** of trypanosomatid parasites from honey bees (*Crithidia mellificae, Lotmaria passim*), bumble bees (*Crithidia bombi*) and mosquitoes (*Crithidia *fasciculata*). The right-most facet label indicates the strain's host of origin. Each point represents the specific growth rate ($h^{-1}$) from one sample. The experiment was conducted over two experimental blocks (Round 1: circles; Round 2: triangles). Lines and shaded bands show predictions and 95% bootstrap confidence intervals from biphasic logistic models. Data from *C. bombi* are taken from [5].

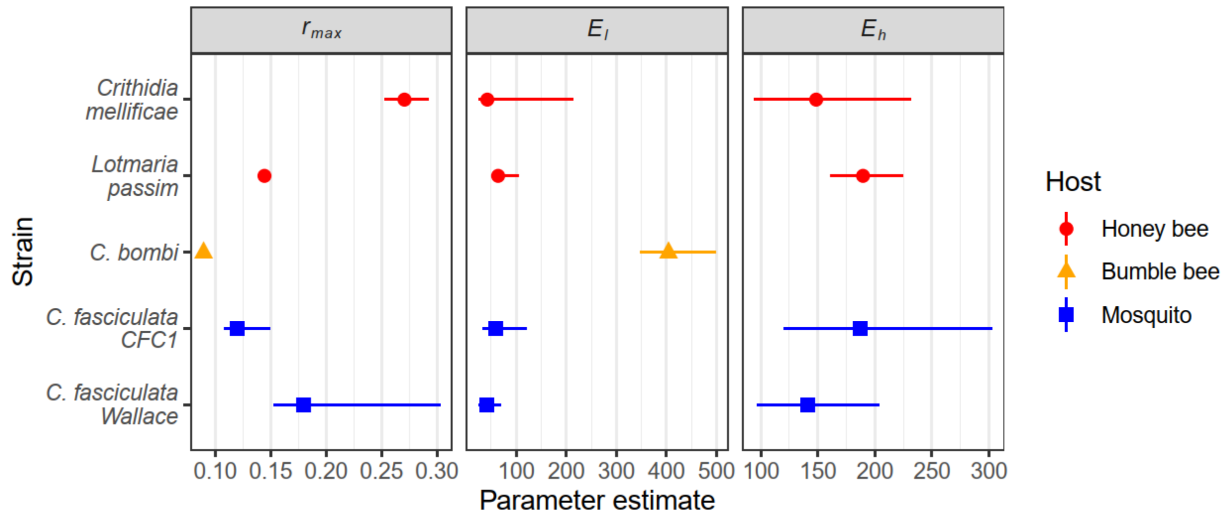

**Supplementary Figure 5. Estimates for rates of peak growth (*$r_{max}$*, in $h^{-1}$) and inhibition due to low (*$E_l$*) and high (*$E_h$*) pH** for parasites of honey bees (*Crithidia mellificae, Lotmaria passim*), bumble bees (*C. bombi* strain IL13.2, tested in [5] for low-pH inhibition only), and mosquitoes (*C. fasciculata*). Points and error bars show estimates and 95% bootstrap confidence intervals for predictions from biphasic logistic models. Colors and shapes correspond to host of origin. See Supplementary Figure 4 for full model predictions for *C. bombi.*

### SUPPLEMENTARY DATA

**Supplementary Data S1:** Growth rates from temperature and pH experiments. Data for *C. bombi* are from our previously published work [4,5].
